# Integrating nested species distribution models and multicriteria prioritization into an open-access geoportal for tropical forest restoration

**DOI:** 10.64898/2026.10.08.757788

**Authors:** Robinson J. Herrera-Feijoo, Jennifer Morales-Barbero, Bolier Torres, Ruben G. Mateo

## Abstract

Tropical forests face increasing pressures from climate change and deforestation, which calls for the selection of climate-resilient species for ecological restoration. Species distribution models (SDMs) calibrated solely with regional data in the tropics often suffer from niche truncation due to data scarcity. We applied spatially nested models (N-SDMs), in one of their first applications in the tropics, to assess the current and future climatic suitability of 199 native forest species of continental Ecuador, integrating global and regional data through a nested approach that consistently outperformed regional-only SDMs (AUC = 0.95 vs. 0.83). Given a widespread projected loss of climatic suitability and an insufficient representation within the National System of Protected Areas, passive conservation alone is insufficient, which calls for an active prioritization tool. We developed a restoration potential index (IPR) that provides a continuous score for each species in every cell by integrating current potential presence, future climatic confidence and documented ecosystem services. This species- and cell-level resolution yields a restoration priority ranking that is robust across climate scenarios and is available for any site in the territory, distinguishing species of consolidated priority from those requiring urgent active propagation owing to their reduced current potential extent. All results are made available through the interactive geoportal geoSABINA, providing the first spatially explicit and climate-informed infrastructure to support the implementation of the Ecuadorian National Forest Restoration Plan (PNRF) and equivalent frameworks in the American tropics.

**HIGHLIGHTS:** Nested SDMs outperformed regional-only SDMs for 199 native tree species of Ecuador.

Most species are projected to lose climatic suitability by 2050, mainly under SSP585.

Protected areas cover <25% of the potential range of 86% of the species.

IPR integrates current presence, future climate suitability and local community uses.

geoSABINA makes the cell-level species ranking available for any site in Ecuador.

## 1. INTRODUCTION

Tropical forests of the Neotropics constitute one of the main global hotspots of plant biodiversity (De Freitas & Junior, 2025; Raven et al., 2020), with Ecuador being a clear reflection of this concentration. A single hectare in Yasuní National Park can hold more tree species than the entire native tree flora of the United States and Canada (Bass et al., 2010). However, this richness faces increasing vulnerability owing to the limited adaptive capacity of tropical flora to climate change (Aguirre-Gutiérrez et al., 2025) and the constant pressure of deforestation and habitat loss (Zhang et al., 2024; Zou et al., 2025). To mitigate these threats, global initiatives such as the Bonn Challenge and the Kunming-Montreal Global Biodiversity Framework promote the large-scale restoration of degraded ecosystems, partly as a response to climate change itself (Gittleman et al., 2026; Hutchinson et al., 2026). Their long-term success depends on selecting, for each site, species suited to its future climate and to the needs of local communities, yet tools integrating both criteria at the scale at which restoration decisions are made are still lacking (Di Sacco et al., 2021).

Species distribution models (SDMs) have become a fundamental tool for assessing risks and informing forest planning (Franklin, 2023; Guisan et al., 2017; Qiao & Escobar, 2025; Thuiller, 2024; Zurell et al., 2023). However, their application in the Neotropics faces severe methodological constraints (Mateo et al., 2010). Occurrence records are scarce and show a strong sampling bias towards accessible areas (Engemann et al., 2015; Enquist et al., 2019; Hughes et al., 2025). Moreover, models are often calibrated only within the study region, frequently because high-resolution covariates (e.g., microtopography or soil properties) are available only at that scale. This truncates the environmental range captured by the model (Chevalier et al., 2022; Goicolea et al., 2025; Guisan et al., 2025), so the modelled niche is incomplete, producing biased predictions and unstable projections with low transferability under future climate scenarios (Brodie et al., 2022; Guisan et al., 2025). To overcome this methodological challenge, spatially nested species distribution models (N-SDMs) hierarchically integrate a global model (which captures the species’ response across its entire distribution, an approximation of its full climatic niche) to constrain and refine the predictions of a regional model (Adde, Rey, Brun, et al., 2023; Guisan et al., 2025; Mateo et al., 2024). Their application, however, has been concentrated in temperate ecosystems, and their extension to the Neotropics remains exceptional (Banks et al., 2026; Dogan & Yilmaz, 2026; Goicolea et al., 2025, 2026).

In this context, continental Ecuador represents an ideal study setting for planning climate-informed restoration interventions. With ∼18,800 vascular plant species, about 4,500 of them endemic (Freire-Fierro et al., 2024; Neill, 2012), the country is experiencing accelerated habitat degradation and loss driven by deforestation, agricultural expansion and extractive activities (Aragon-Osejo et al., 2025; Cogălniceanu & Szekely, 2025), reducing its capacity for natural regeneration (MAE, 2023). Given that a static network of protected areas alone does not guarantee the long-term persistence of biodiversity in the face of the pace of climate change (Fajardo et al., 2023; Li et al., 2024; Watson et al., 2023), it is essential to combine passive conservation with active restoration planned for specific sites. Ecuador has identified approximately 4.54 million ha with potential for forest restoration under the National Forest Restoration Plan 2019–2030 (MAE, 2019), while the emerging national restoration framework for 2026–2035 proposes bringing approximately 150,000 ha under restoration by 2035 (MAE et al., 2026). The substantial gap between the extent with restoration potential and the area that can realistically be addressed underscores the need for spatially explicit prioritization. Such prioritization should determine not only where restoration efforts are most strategic and which species are most likely to persist under future climatic conditions, but also which species can provide ecosystem services and contribute to local livelihoods, thereby strengthening the ecological and social sustainability of restoration investments (Krainovic et al., 2025; Martin et al., 2021; Torres et al., 2023).

Although some restoration initiatives, particularly in Europe, have begun to incorporate future climate suitability into species selection (Carrillo-García et al., 2023; Gastón et al., 2014), restoration species have traditionally been selected for their current climatic suitability or their commercial value alone, without considering whether they will remain suitable under future climate or whether they meet the social and cultural demands that determine local acceptance of restoration. This compromises the long-term sustainability of the intervention (Barragán et al., 2023; Barragán & Rhemtulla, 2025; Fremout et al., 2022; Hernandez-Montilla et al., 2026). In addition, the scientific knowledge generated rarely reaches managers in a usable format, as the outputs of predictive models usually remain confined to static maps in the supplementary material of publications. To overcome this operational limitation, interactive open-access geoportals are emerging in Europe as a key infrastructure that makes such results available to decision-makers in a dynamic format directly applicable to territorial planning (Chacón-Moreno et al., 2026; Goicolea et al., 2026).

In this study, we developed and applied a comprehensive forest restoration framework for 199 native tree species of continental Ecuador, combining, for the first time in a Tropical country, spatially nested models (N-SDMs), a multicriteria prioritization index (IPR) and an open-access geoportal (geoSABINA), with the aim of providing a spatially explicit, climatically robust and socially relevant basis to guide the transition from passive conservation to active restoration in national strategies. To this end, this work addresses three specific objectives: (i) to assess the ability of N-SDMs to mitigate niche truncation and improve predictive stability compared with traditional regional SDMs (hereafter regional-only SDMs); (ii) to characterize the spatial patterns of potential richness, their dynamics under climate change scenarios and the representativeness gap of the National System of Protected Areas (NSPA), as a baseline diagnosis for prioritization; and (iii) to prioritize species for each site through a multidimensional Restoration Potential Index (IPR), calculated for each species in every cell, that integrates current potential presence, future climatic confidence and the ecosystem services of interest to local communities, making these results available through the interactive geoportal geoSABINA. Together, this methodological framework consolidates the first spatially explicit and dynamic infrastructure to guide the implementation and updating of the PNRF and equivalent frameworks in the American tropics.

## 2. MATERIAL AND METHODS

### 2.1. Species and occurrence records

We modelled the current and future distribution of 199 native forest species of socioeconomic and ecological relevance in Ecuador, selected from the Forest Administration System (López-Tobar et al., 2023, 2024, 2026). To avoid ecological niche truncation and the sampling biases characteristic of tropical regions, two spatial extents were defined at a common spatial resolution of 1 km. At the global scale (the species’ broad range), the calibration area extended from northern Canada to southern Argentina (∼104.5 million km²), using occurrence records retrieved from the Global Biodiversity Information Facility (GBIF) and the Botanical Information and Ecology Network (BIEN) database (Enquist et al., 2026). At the regional scale (the study area), calibration was restricted to continental Ecuador (∼248,500 km²), using records from the Biodiversity Information System of Ecuador (SISBIO; https://bndb.sisbioecuador.bio/bndb/).

Occurrence records were cleaned following a spatial and taxonomic protocol (Kass et al., 2025; Moudrý et al., 2024). Only botanical specimens collected after 1950 were retained, ensuring taxonomic reliability and temporal consistency with the climatic covariates (Daru, 2025; Humphreys et al., 2025; Moudrý et al., 2024). Imprecise coordinates and duplicate records were removed with *CoordinateCleaner* (Zizka et al., 2019), and the remaining records were thinned to one per 1 km² cell with *GeoThinneR* (Mestre-Tomás, 2025) to reduce sampling bias. Finally, only species with at least 15 independent records at both calibration scales were retained, avoiding biases associated with small sample sizes (Collart & Guisan, 2023; Gaul et al., 2020).

### 2.2. Covariates and scenarios

At the global scale, model calibration was based exclusively on bioclimatic covariates. At the regional scale, the same set of covariates was adopted after verifying, through a preliminary exploratory analysis in continental Ecuador, that the inclusion of edaphic and/or topographic variables did not provide consistent improvements in predictive performance compared with a purely climatic model, for both regional-only SDMs and N-SDMs (details in Supplementary Material S1.1).

The current baseline climate was characterized using bioclimatic covariates obtained from the WorldClim v2.1 database (reference period 1970–2000, 30 arc-second spatial resolution, ∼1 km) (Fick & Hijmans, 2017). To avoid the artificial spatial discontinuities generated by quarter-based predictors in equatorial regions (Booth, 2022), BIO8, BIO9, BIO18 and BIO19 were excluded, retaining a final set of 15 bioclimatic covariates.

Future climate was characterized using the same 15 bioclimatic covariates from WorldClim v2.1 (period 2041–2060, hereafter 2050; 30 arc-second spatial resolution, ∼1 km), derived from CMIP6 (Tebaldi et al., 2021). To account for the climatic uncertainty inherent in modelling, four general circulation models were selected (GCMs: ACCESS-CM2, GISS-E2-1-G, HadGEM3-GC31-LL, MIROC6), and future suitability was assessed under two socioeconomic pathways: a strong mitigation scenario (SSP126) and a very high emissions scenario without active mitigation (SSP585) (Meinshausen et al., 2020; Tebaldi et al., 2021).

### 2.3. Spatially nested modelling (N-SDMs) and regional-only SDMs

We used the sabinaNSDM package (Mateo et al., 2024) to estimate species’ climatic suitability, comparing two modelling strategies: a regional-only SDM, calibrated solely with regional-scale records and bioclimatic covariates, versus a spatially nested model (N-SDM) based on the *covariate* strategy (see Mateo et al, 2024, Guisan et al. 2025), in which the suitability prediction of the global model is incorporated as an additional covariate in the regional model (Fig. 1).

**Figure 1.**
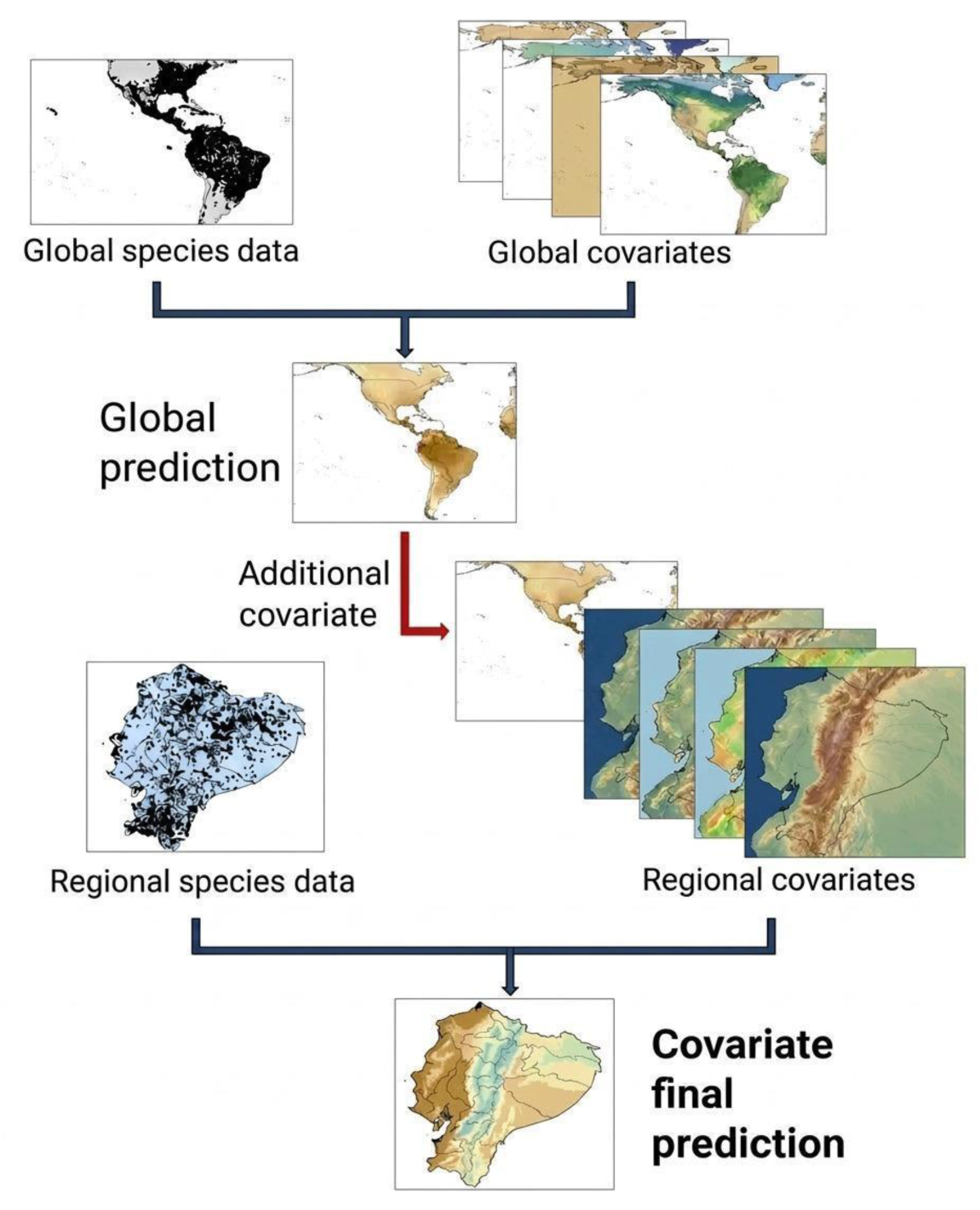
Conceptual framework of the *covariate* strategy within the spatially nested modelling approach (N-SDM). Global-scale data and covariates generate a global prediction, which is incorporated as an additional covariate together with the regional data and covariates to produce the final prediction for continental Ecuador.

Covariates were selected with the automated two-step embedded procedure of sabinaNSDM, which first removes collinear covariates using pairwise Pearson correlations (|r| < 0.70) and then ranks and retains the most informative ones through regularization (Adde, Rey, Fopp, et al., 2023).

Distributions were estimated through an ensemble forecasting approach (Araujo & New, 2007; Marmion et al., 2009; Thuiller et al., 2009), combining three algorithms that differ in how they model species–environment relationships: generalized additive models (GAM; Hastie & Hastie, 1991), boosted regression trees (BRT; Ke et al., 2017) and random forests (RF; Breiman, 2001). As true absences were unavailable, 10,000 random pseudo-absences were generated (excluding cells with known occurrences), weighted to a prevalence of 0.5 to balance the contribution of both classes during model fitting and avoid imbalance-related biases (Barbet-Massin et al., 2012). For each algorithm, data were split into training (80%) and evaluation (20%) sets in ten random replicates (Moudrý et al., 2024), yielding 30 calibrated models per species and strategy, which were then projected onto current conditions and each of the eight future scenarios. Performance was evaluated using the area under the ROC curve (AUC) (Fielding & Bell, 1997), the True Skill Statistic (TSS) (Allouche et al., 2006) and the Continuous Boyce Index (CBI) (Hirzel et al., 2006).

Replicates with AUC < 0.80 were discarded (Herrera-Feijoo et al., 2023; Mateo et al., 2013), and the remaining ones were combined into a consensus model as an AUC-weighted mean, so that better-performing replicates contribute more to the final prediction (Marmion et al., 2009). The final consensus models of both strategies (regional-only SDM and N-SDM) were projected onto the current climate and the eight future scenarios (4 GCMs × 2 SSPs). These continuous predictions were binarized using the threshold that jointly maximizes sensitivity and specificity on the ROC curve (Hellegers et al., 2025; Scherrer et al., 2018), yielding binary potential presence–absence maps (1/0).

### 2.4. Potential biodiversity

Potential species richness was estimated by cell-by-cell stacking of the current and future binary potential presence–absence maps (Stack-SDMs, Mateo et al., 2012). Changes at the species level were assessed by comparing the current and future binary potential presence–absence maps under each scenario. A cell was considered suitable in the future when at least one of the four GCMs predicted presence. This inclusive criterion captures the full range of future conditions projected by the GCMs, so losses should be read as minimum estimates, and stability and gain as maximum estimates (Porfirio et al., 2014). Agreement among GCMs is accounted for separately in the IPR through future climatic confidence. Each cell was then classified into three categories: (i) stability (current and future potential presence), (ii) loss (potential presence restricted to the present) or (iii) habitat gain (current absence and future presence).

### 2.5. Representativeness of the NSPA

We evaluated the representativeness of Ecuador’s National System of Protected Areas (NSPA) through a conservation gap analysis, using the official boundaries from the Spatial Data Infrastructure (IDE) of the Ministry of the Environment of Ecuador (http://ide.ambiente.gob.ec/mapainteractivo/). Current binary potential presence–absence maps were spatially overlaid with the official NSPA cartography to calculate the percentage of each species’ potential presence area contained within the network, considering a cell as protected when its centroid fell within the NSPA boundaries.

### 2.6. Restoration Potential Index (IPR)

To identify and rank the species with the highest restoration potential, we developed a Restoration Potential Index (IPR), calculated for each species in every cell under the SSP126 and SSP585 scenarios. The index multiplicatively integrates three components: 1) the current potential presence of the species in the cell, 2) future climatic confidence (long-term persistence, assessed as the proportion of GCMs agreeing in predicting presence) and 3) the ecosystem services provided by the species.

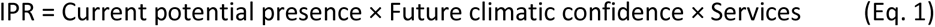

Current potential presence corresponds to the binary potential presence–absence map of each species. Climatic confidence provides a continuous measure (0, 0.25, 0.50, 0.75, 1.00) of the agreement among the four GCMs considered. The Services value of each species was obtained as the simple arithmetic mean of seven variables standardized to a common 0–1 scale. Specifically, the effectively harvested timber volume (VP), obtained from the official records of the Forest Administration System of Ecuador (https://saf.ambiente.gob.ec) and recent literature (López-Tobar et al., 2024, 2026), showed a strongly skewed distribution and was therefore log-transformed (log10) prior to min–max scaling. Wood density (WB) was extracted from the Global Wood Density Database, which compiles 16,468 wood density records for 8,412 taxa worldwide (Chave et al., 2009), using the *BIOMASS* package (Réjou-Méchain et al., 2017), employing regional estimates (*SouthAmericaTrop*) when species-specific data were unavailable and treating density values equal to 0 as missing data. The remaining five variables [timber use (UM; timber/non-timber), food for humans (FH), food for animals (FA), medicinal use (MU) and agroforestry support (SAF)] were coded as binary (1 = documented use, 0 = no evidence) from the Useful Plants of Ecuador database (de la Torre, Lucía et al., 2008), the outcome of five years of fieldwork and the review of ∼500,000 herbarium specimens (with 19,611 records documenting uses for 5,172 species). The absence of a record (value 0) indicates undocumented use (Data S2) and not necessarily non-existent use. The resulting IPR values are bounded between 0 and 1. A value of 0 in any of the three components nullifies the index for that species in that cell.

The mean IPR of each cell was calculated as the arithmetic mean of the IPR of all species present. This cartographic synthesis complements the individual species values to show the overall restoration priority. As a robustness check against the instability of this average in low-richness cells, a corrected version of the mean IPR was additionally calculated using an empirical Bayes shrinkage estimator (further details in Supplementary Material S1.2).

To evaluate the robustness of the IPR prioritization, its sensitivity was analyzed using a leave-one-out approach, excluding each of the seven Services variables in turn. This matters because five of these variables come from a single source of binary data (de la Torre et al., 2008), which does not distinguish between undocumented and non-existent use, nor does it provide measures of completeness or uncertainty associated with each species or region. In each cell with at least five species, the species ranking obtained with all variables was compared with the ranking obtained after removing each variable, using three metrics, and the values were then averaged across cells: Spearman’s coefficient (agreement of the complete species ranking), the overlap among the five highest-priority species (top-5) and the Rank-Biased Overlap (RBO; (Webber et al., 2010), which gives particular weight to agreement in the top positions of the ranking (further details in Supplementary Material S1.3).

To summarize the IPR at the species level, each species was classified by combining its mean IPR across its current potential distribution with its current potential extent (number of cells), both split at their median, separately for each scenario (further details in Supplementary Material S1.4). Species without a defined IPR under a given scenario, that is, whose current range does not overlap with any area predicted as suitable by at least one GCM, were not included in this classification.

### 2.7. Interactive geoportal

All results were made available through geoSABINA, a free-access interactive geoportal (https://www.geosabina.com/ecuador-geoportal/). For each forest species assessed, this interactive web platform allows users to visualize, explore and download the current climatic suitability; the future projections corresponding to the eight combinations resulting from crossing the 4 GCMs and the 2 SSPs for 2050; and, where the IPR is defined, its value for each species in each cell. By selecting any site in continental Ecuador and a scenario, users obtain the list of species ranked by their IPR and climatic suitability.

## 3. RESULTS

### 3.1. N-SDMs vs regional-only SDMs

N-SDMs achieved the highest predictive performance (AUC = 0.95 ± 0.05; TSS = 0.61 ± 0.30; CBI = 0.61 ± 0.29), consistently outperforming regional-only SDMs (AUC = 0.83 ± 0.11; TSS = 0.38 ± 0.29; CBI = 0.54 ± 0.31). Likewise, N-SDMs reduced the standard deviation of AUC across species, which increases the stability and certainty of projections compared with regional-only SDMs. According to the exploratory analysis detailed in Supplementary Material S1.1, adding edaphic and topographic covariates did not improve predictive performance over bioclimatic covariates alone, so they were discarded for parsimony and to avoid overfitting and overparameterization (Moreno-Amat et al., 2015). Species-level performance metrics are available in the Supplementary Material (Data S1).

### 3.2. Spatio-temporal patterns of potential biodiversity

Under current climatic conditions, the stacked-SDMs derived from N-SDMs showed a marked concentration of species towards the central-eastern and central-southern Ecuadorian Amazon, identifying Morona Santiago province as the main hotspot of potential richness in the continental territory, with a maximum of 137 species per cell across the whole study area. Potential richness decreases progressively towards the Andes, where it is substantially reduced in high-Andean environments, and towards the Coast (Fig. 2). Future loss of suitability is concentrated in the central-northern Amazon. Gain areas are concentrated in the Andean– Amazonian foothills in both the north and the south, whereas stability is more diffusely distributed, mainly in the Andean and western transition provinces.

**Figure 2.**
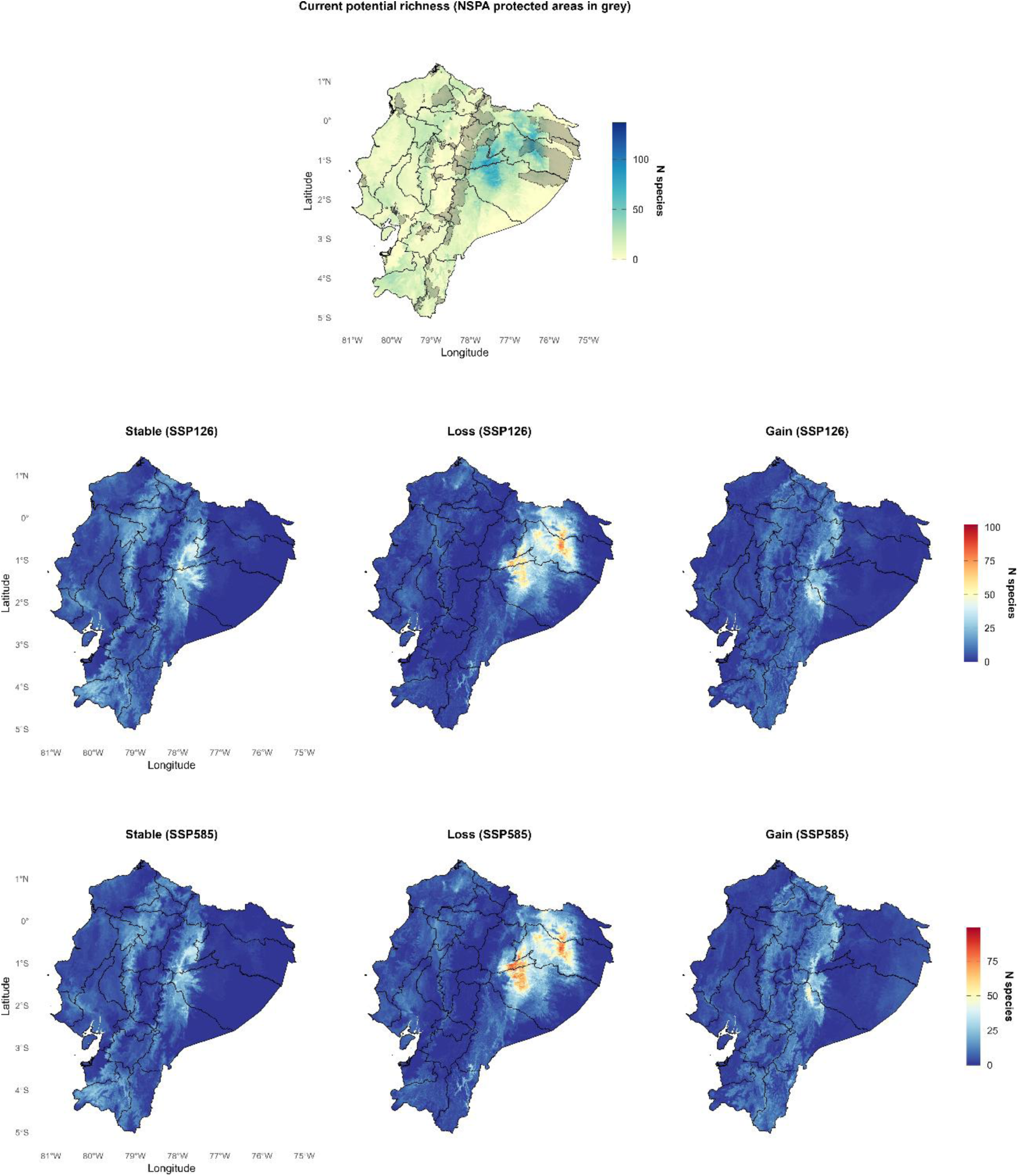
Current potential richness (NSPA protected areas in grey) and species-level habitat stability, loss and gain under SSP126 and SSP585. Future presence was defined with an inclusive criterion (≥1 of 4 GCMs in agreement). Values show the number of species per cell per category.

The magnitude of projected change was heterogeneous among species, but the loss of suitable potential habitat clearly dominated over gain in both scenarios. The median species lost 59.0% of its current potential distribution under SSP126 and 70.5% under SSP585. Under SSP126, 32.66% of species retained less than a quarter of their current area as climatically stable (i.e., they lost more than three quarters of their distribution), a proportion that rose to 46.23% under SSP585. Climatic gains were, in contrast, much more limited. In total, 35.18% of species under SSP126 and 40.70% under SSP585 gained less than a quarter of their current extent in new areas of suitable habitat. The more severe scenario thus aggravated the magnitude of loss without proportionally expanding the availability of new suitable habitat (Data S1). This heterogeneity among species is precisely what the IPR translates into an operational cell-level prioritization.

### 3.3. Conservation gaps in the NSPA

The NSPA poorly represents the potential distribution of most species, as 86.43% of them had less than a quarter (< 25%) of their potential distribution within the boundaries of the protected system (Fig. 2; Data S1).

### 3.4. Species prioritization for restoration

The mean IPR (Fig. 3) showed a consistent spatial pattern across both scenarios, with the highest values in the Andean region, its foothills and the transition zones towards the Coast and the Amazon, and the lowest values in the lowlands of the central-eastern Amazon. This pattern held after applying the empirical Bayes shrinkage correction for the instability of the average in low-richness cells [k = 5.77 (SSP126) and 5.76 (SSP585); Supplementary Material S1.2]. This map illustrates the regional pattern of the index. The prioritization with direct application to decision-making is presented below at the species and cell level, and is available interactively in the geoportal.

**Figure 3.**
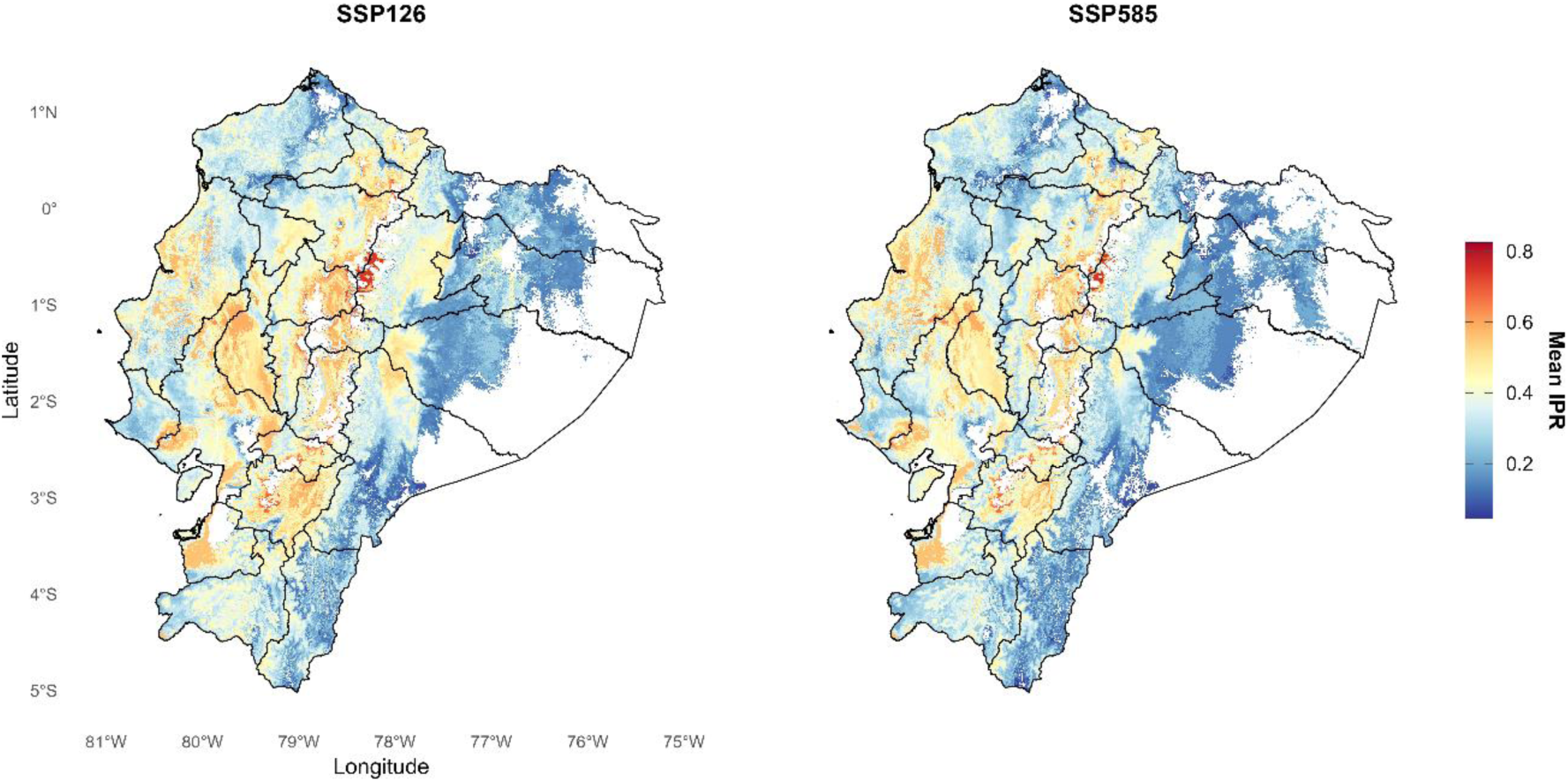
Mean Restoration Potential Index (IPR) per cell under SSP126 and SSP585, integrating current and future climatic suitability with the Services dimension.

The prioritization was robust to the exclusion of individual ecosystem services. When each was excluded and the IPR recalculated, the within-cell species ranking maintained a high Spearman correlation (ρ = 0.93–0.98), a high overlap among the five highest-priority species (top-5 = 90– 97%) and a moderate-to-high rank-weighted overlap (RBO, which prioritizes the top positions of the ranking = 0.57–0.64) relative to the full ranking, across the seven variables evaluated. Food for humans (FH), medicinal use (MU) and food for animals (FA) showed the greatest relative influence on all three metrics, with values very close to one another (minimal differences that cannot be ranked with confidence). No individual service alone determines the final prioritization (details by service and metric in Supplementary Material S1.3).

Of the 199 species modelled, the IPR was undefined for 13 (6.5%) under SSP585 —4 of which also under SSP126— as there was no cell where their current distribution spatially overlapped with any area predicted as suitable by at least one GCM. Of the 186 species with a computable IPR under both scenarios, 59 (31.72%) remained consistently in the highest restoration priority quadrant (high IPR–high extent; Fig. 4), with an overall classification agreement between scenarios of 90.32%, and a second group of 27 species (14.52%) remained in the high IPR–low extent quadrant, combining high future climatic confidence with a small current potential extent. A third group of 28 species (15.05%) remained in the low IPR–high extent quadrant.

**Figure 4.**
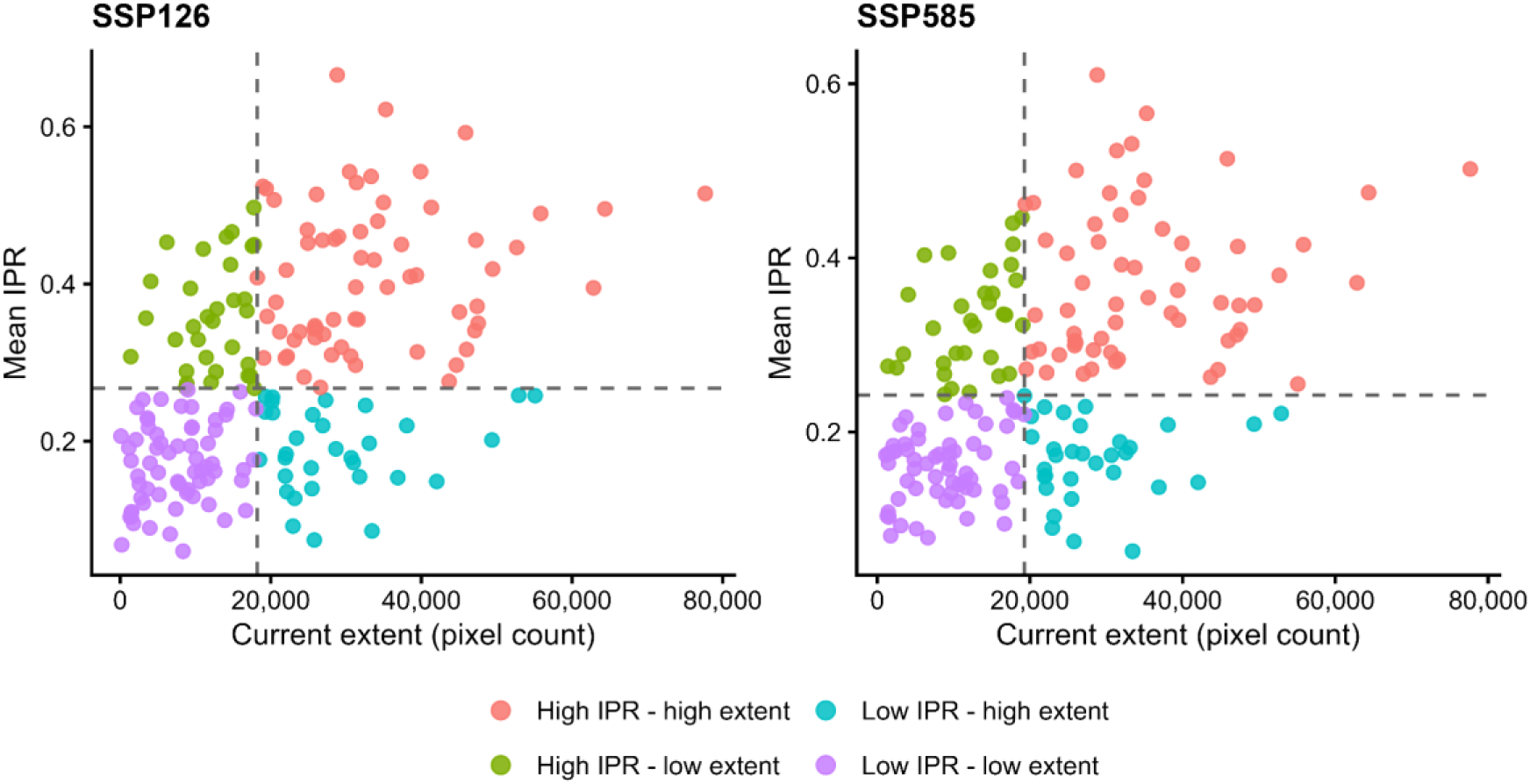
Species-level restoration potential, mean IPR versus current potential extent (number of suitable cells). Dashed lines mark the median IPR and median extent across species; species fall into four groups, high IPR–high extent, high IPR–low extent, low IPR–high extent, and low IPR–low extent.

As a concrete example of this cell-level prioritization capacity, the cell with the highest potential richness (Morona Santiago) also concentrates 20 of these 27 priority species, the highest co-occurrence of the group across the entire continental territory. These species share a predominantly timber-use profile (present in all of them), combined in most cases with medicinal and food uses, and include highly recognized taxa such as *Cinchona officinalis* L. (Ecuador’s national tree) and *Ilex guayusa* Loes.

As a final operational product, the projections, change dynamics and spatial IPR classification of all species were integrated into the open-access geoportal geoSABINA https://www.geosabina.com/ecuador-geoportal/, where the complete ranking can be explored interactively for this or any other cell in the territory (Fig. 5).

**Figure 5.**
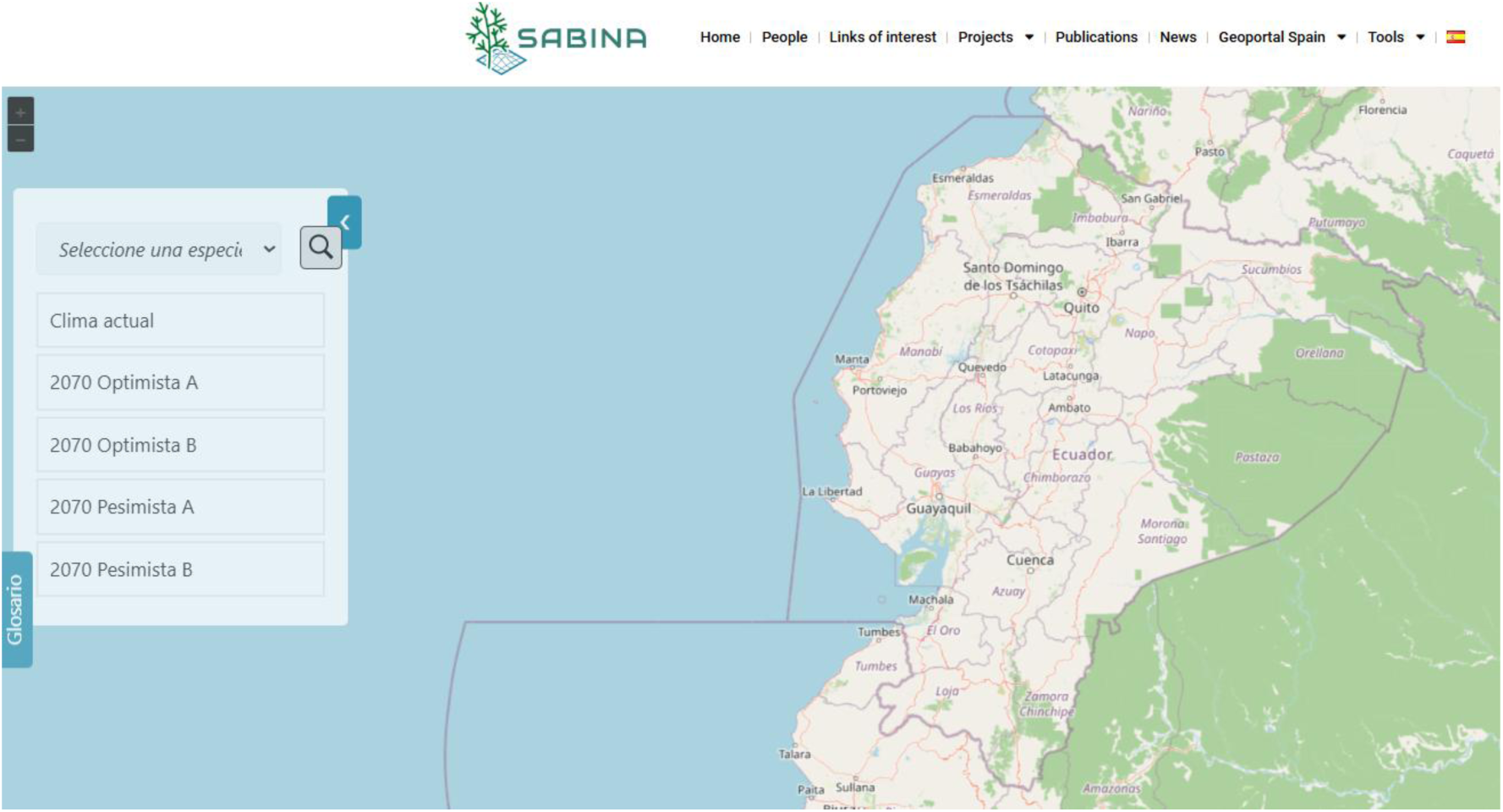
Application case of the geoSABINA geoportal in a representative cell (Morona Santiago, 1.46°S – 78.19°W) that concentrates both the highest potential species richness nationwide (137 species) and 20 of the 27 robustly identified urgent-priority species (high IPR – low extent).

## 4. DISCUSSION

The long-term effectiveness of restoration depends on selecting, for each site, species suited to its future climate and to the needs of local communities. Our results project a widespread loss of climatic suitability for the 199 native tree species analyzed and show that the static protected area network (NSPA) cannot mitigate this impact. Combining N-SDMs, a multicriteria index (IPR) and an open-access geoportal (geoSABINA) provides a dynamic decision-support platform for planning restoration.

### 4.1. Spatially nested models (N-SDMs) in the Neotropics

To our knowledge, this study constitutes one of the first applications of spatially nested distribution models (N-SDMs) in a megadiverse tropical region and with explicit ecological restoration goals. Previous applications of this approach have mainly focused on Europe and other temperate regions, where the high density of biological records, consolidated monitoring networks and the availability of detailed environmental information favor model calibration (Adde et al., 2024; Chevalier et al., 2021, 2022; Goicolea et al., 2025; Scherrer et al., 2018). In contrast, applying this approach in tropical regions such as Ecuador represents a more demanding methodological test, owing to the scarcity and spatial bias of occurrence records, which are frequently concentrated near roads, settlements and easily accessible areas (Engemann et al., 2015; Hughes et al., 2025; Lobos et al., 2026). Despite these limitations, N-SDMs performed well across all evaluation metrics, consistent with the improvement reported for this approach in other contexts (Goicolea et al., 2025; Mateo et al., 2019, 2024).

The conceptual value of this approach in the tropics differs substantially from its role in temperate latitudes. In Europe, niche truncation is mainly an administrative artefact caused by political borders that artificially cut species’ distributions (Chevalier et al., 2022; Goicolea et al., 2025). In the tropics, by contrast, the problem stems from a severe Wallacean shortfall (Engemann et al., 2015; Hughes et al., 2025) and a complex biogeographic history characterized by high rates of endemism and abrupt, complex Andean–Amazonian gradients that hinder the full capture of environmental variation at the local scale. Consequently, the mechanism by which the N-SDM approach mitigates truncation in the tropics operates differently from the European case. Rather than merely “completing” a transboundary distribution range geographically, spatial nesting acts as an ecological stability filter. By incorporating the global prediction as a covariate, the regional model is partially constrained by the niche limits at a broader scale (Guisan et al., 2025), which helps reduce the overfitting arising from local sampling biases, such as the accumulation of records along accessible roads, and favors more stable projections, especially for species with scarce data or wide but fragmented distributions (Anselmetto et al., 2025; Lobos et al., 2026). Uncertainty is also considered at several stages of the workflow. Algorithm uncertainty is addressed through the ensemble of three algorithms and ten replicates, inter-GCM uncertainty through future climatic confidence in the IPR, and emission scenario uncertainty by comparing SSP126 and SSP585.

N-SDMs calibrated exclusively with bioclimatic variables performed as well as those incorporating edaphic and topographic covariates (Supplementary Material S1.1). As these additional predictors did not improve predictive performance, the bioclimatic set was retained for parsimony and to reduce the risk of overfitting (Moreno-Amat et al 2015). This contrasts with findings in temperate regions (Adde et al., 2024; Broennimann & Guisan, 2026; Sanguet et al., 2022) and suggests that climate may represent a dominant environmental filter at the spatial scale and resolution for the species considered in this study. One possible ecological explanation is that, in humid tropical forests, high precipitation and relatively continuous water availability may reduce the influence of topographically driven differences in water availability and evapotranspiration compared with water-limited ecosystems, such as Mediterranean forests, where slope orientation can strongly determine vegetation patterns. However, this explanation may not apply to seasonally dry tropical forests or other environments where water availability is limiting. The limited contribution of edaphic variables may also reflect the ecological characteristics of the species examined and the spatial resolution and accuracy of SoilGrids in tropical regions (Malhotra et al., 2026; Poggio et al., 2021). Similarly, topographic effects operating at finer spatial scales may not be adequately captured at the approximately 1 km resolution used in this study. Therefore, the limited predictive contribution of edaphic and topographic variables should not be generalized to other species, ecosystems or spatial scales, where these factors may play a more important role.

### 4.2. Forest distribution dynamics and restoration

The concentration of current potential richness in the Ecuadorian Amazon and the Andean– Amazonian transition zones, with Morona Santiago province as the main hotspot, is consistent with patterns reported for other taxonomic groups in the region, associated with topographic heterogeneity and altitudinal gradients (Lessmann et al., 2014; Mateo et al., 2012, 2013). This high diversity results from altitudinal gradients that favor the overlap of lowland and montane species niches (Engemann et al., 2015; Malizia et al., 2020) and from the high levels of endemism of montane forests (Balslev, 1988; Richter et al., 2009). The lower potential tree species richness estimated in the Andes, particularly at high elevations such as the páramo ecosystems, and in the Coastal region may reflect physiological constraints (Guzmán et al., 2026; Restrepo et al., 2025) imposed by environmental conditions. However, the relatively low predicted richness of tree species in páramo ecosystems also reflects their natural dominance by non-arboreal vegetation and should not be interpreted as indicating lower biodiversity or conservation value. These ecosystems harbour numerous endemic species, highlighting the importance of conservation strategies that consider not only species richness but also endemism and ecological uniqueness (Mateo et al., 2013).

In the future, climate projections warn of a drastic contraction of suitable habitat in the central-northern Amazon (Sucumbíos and Orellana provinces), where the loss of climatic suitability is concentrated (Fig. 2). Areas of climatic gain, although smaller in magnitude than those of loss, are concentrated in the Andean–Amazonian foothills in both the north and the south, whereas the few stable areas, located mainly in the Andean and western transition provinces, may act as climatic refugia. Their high topoclimatic heterogeneity may favor the persistence of suitable conditions and the formation of microrefugia (Fadrique et al., 2026; Guzmán et al., 2026). Focusing restoration on these transition zones, rather than applying uniform strategies across the Amazonian lowlands, could therefore favor species persistence, the provision of ecosystem services and landscape resilience (Rosenfield et al., 2025; Song et al., 2026). Given that natural colonization is unlikely to keep pace with climate change, constrained by slow forest dynamics and the dispersal capacity of each species (Farfan-Rios et al., 2025; Feeley et al., 2023; Xu et al., 2025), these areas should be prioritized for active restoration, where assisted planting can anticipate the colonization that natural processes would not achieve in time. Consequently, active restoration programs must overcome the limitations of static protected areas by prioritizing climatically resilient, multipurpose species that strengthen ecosystem functionality and local livelihoods, while conserving refugia and enhancing connectivity between stable areas and gain zones outside the NSPA.

Importantly, these spatial priorities should not be interpreted as implying a uniform tree-planting strategy. In Ecuador, restoration occurs across a mosaic of degraded forests, pastures, agricultural lands and agroforestry systems, requiring different intervention pathways ranging from assisted natural regeneration and enrichment planting to mixed-species plantations, agroforestry and silvopastoral systems. The species-level climatic information provided here can therefore complement land-use and degradation criteria to determine not only where restoration is needed, but which species assemblages are more appropriate for each restoration pathway.

### 4.3. The IPR as a multicriteria prioritization tool

Forest restoration can simultaneously contribute to ecological, productive and social objectives (Brancalion et al., 2025; Owusu Danquah et al., 2026). However, species selection has traditionally relied on known environmental tolerances, local site conditions or socioeconomic value, often without integrating these criteria or accounting for future climate change. Complementing this information with current and future climatic suitability predictions can support a transition from static to dynamic, climate-adaptive restoration planning (Carrillo-García et al., 2023). Moreover, the marked interspecific variation in climate change responses observed in our study highlights the importance of integrating climatic persistence and socioeconomic value into species-specific restoration priorities (Barragán & Rhemtulla, 2025; Fremout et al., 2020; García Del Barrio et al., 2026).

The IPR directly addresses this limitation by integrating, multiplicatively and cell by cell, current potential presence, future climatic confidence (persistence of suitable habitat) and a ecosystem services dimension that simultaneously combines productive relevance (wood density, effectively harvested timber volume) and documented uses of local value (timber, medicinal, food, agroforestry support) (de la Torre et al., 2008; López-Tobar et al., 2024, 2026; Réjou-Méchain et al., 2017). Our approach thus enables a balanced selection, preventing a restoration program from relying exclusively on commercially attractive taxa with high future climatic vulnerability, and likewise preventing the selection of climatically stable species of little value for local livelihoods (Barragán et al., 2023). As the ecosystem services dimension averages several variables, the index tends to favor species with multiple documented uses over those with a single high-value use, which managers should consider when interpreting the ranking. However, the usefulness of the IPR depends not only on identifying priority species and areas but also on the robustness of the resulting priorities, since selecting species and sites based on biased or highly variable estimates may lead to unreliable decisions and investments with poor ecological performance(Brancalion & Holl, 2020; Erbaugh et al., 2020; Marshall et al., 2023). This is particularly relevant here, given that five of the seven ecosystem services variables come from a single binary data source, although the sensitivity analysis showed that no individual variable determines the ranking.

The quadrant scheme (Fig. 4) translates the IPR into four management profiles, and most species kept the same profile under both scenarios, indicating that their priority does not depend on the emission scenario considered. High IPR–high extent species are immediate candidates for large-scale restoration, although local provenance and genetic diversity criteria remain necessary for their adaptive persistence (Barragán & Rhemtulla, 2025; Erickson & Halford, 2020; Guevara et al., 2026; Thomas et al., 2014). High IPR–low extent species should remain a priority despite their scarcity, directing investment towards nursery production and propagation before planting(Andres et al., 2024; Edwards & Cerullo, 2024; Holl et al., 2022; Lacoretz & Pérez, 2025; Thirisha Rani et al., 2026) and preventing restoration from relying on a few easily accessible generalist species. Low IPR–high extent species add little as main restoration species, but they can support the establishment of priority species, for example as facilitators (Galindo et al., 2017; Lortie et al., 2022). Finally, low IPR–low extent species are, as a group, of low cost-effectiveness for restoration (Neeson et al., 2018), although their functional uniqueness or local cultural value, not captured by the IPR, may justify a case-by-case evaluation (Jensen et al., 2021; Mouillot et al., 2013). While this classification summarizes priorities at the species level, the geoportal applies the same logic to each cell, providing a species ranking for any site (Fig. 5). Thirteen species (6.5% of the total modelled) lack an IPR value under SSP585, four of which also under SSP126. Their current distribution range does not share any cell with an area predicted as suitable by at least one GCM, so no site can both host the species today and remain climatically suitable in 2050. For this group, in situ restoration is no longer a viable option, regardless of the ecological or productive relevance of these species, because the IPR cannot recommend where to plant them if no such site exists. Their conservation requires measures beyond the PNRF, such as seed banks, ex situ living collections or assisted translocation towards the areas of climatic gain identified here, which a spatial prioritization index cannot address.

This index can be applied at different planning scales, from the PNRF at the national level to specific territorial initiatives such as the Amazon Biocorridor Program (FBA, 2026), which directs investment towards ecological connectivity in the Ecuadorian Amazon. By directing territorial investment towards the NSPA gaps, the IPR directly supports the implementation of the PNRF and contributes to the functional connectivity of ecological corridors under climate change scenarios.

This multifunctional dimension is particularly relevant in biocultural productive landscapes, where restoration must coexist with farming systems, culturally embedded management and household livelihood strategies (Torres et al., 2026). In these landscapes, native trees are not merely components of ecological recovery but multifunctional assets intentionally retained or established for timber, food, medicinal, cultural and productive functions. Species combining these livelihood functions with high projected climatic persistence could therefore support restoration pathways across agroforestry and silvopastoral systems, live fences, riparian buffers and enrichment plantings (Torres et al., 2023). By integrating climatic persistence with species multifunctionality, the IPR can help identify species portfolios that simultaneously strengthen biodiversity, ecosystem resilience and local livelihoods, extending climate-informed restoration beyond protected forests into the productive landscapes where people live and manage trees.

### 4.4. The geoSABINA geoportal as a dynamic infrastructure for restoration

There are notable precedents of spatial platforms, such as BioModelos in Colombia for expert validation of distribution maps (Velásquez-Tibatá et al., 2019), or climatic suitability based on SDMs viewers developed in Spain (Chacón-Moreno et al., 2026; Goicolea et al., 2026). However, to our knowledge, no equivalent infrastructure exists for climate-informed forest restoration in the American tropics that extends this climatic basis by integrating a multidimensional prioritization such as the IPR.

In contrast to traditional static maps relegated to the supplementary material of publications, the interactive geoportal (https://www.geosabina.com/ecuador-geoportal/) operates as the main knowledge transfer mechanism of the study through an open, persistent and updatable scientific infrastructure that connects directly with the open science movement. Its practical value lies in converting the models for 199 forest species and their projections under multiple future scenarios into information accessible to the Ministry of Environment and Energy, PNRF managers, Decentralized Autonomous Governments (GADs), nursery growers and community organizations, without requiring advanced modelling expertise. By providing easy access to climatic suitability and IPR-based priorities, the geoportal helps decide which species to use in each region, where to intervene, which species to produce in nurseries and how to allocate resources to species and sites that combine climatic persistence, ecosystem functionality and local livelihoods. Providing local actors and communities with these tools helps close the gap between science and decision-making in forest restoration.

This platform provides spatially explicit and dynamic support for species prioritization and selection in forest restoration strategies, complementing the areas identified under the PNRF 2019–2030 (MAE, 2019) and providing an operational basis for implementing the emerging National Restoration Plan 2026–2035 (MAE et al., 2026). By identifying which native species are most likely to remain climatically suitable at a given location, geoSABINA can translate national restoration priorities into site-specific, climate-informed species portfolios.

## 5. CONCLUSIONS

This study presents the first comprehensive species prioritization framework for climate-informed forest restoration in a megadiverse tropical country, combining spatially nested models, an explicit cell-level multicriteria index and a digital knowledge transfer infrastructure. The projected loss of climatic suitability is widespread and far exceeds gains, especially under SSP585, and most species are poorly represented in the NSPA, which underlines the importance of guiding active restoration in degraded areas with species suited to future conditions. The IPR provides a continuous score for each species in every cell, generating a priority ranking that can be applied to any area of interest. This ranking distinguishes species of consolidated priority from those that, despite their high future suitability, require active propagation because of their reduced current potential extent. It also identifies a small group of species (6.5%) whose current and future ranges do not overlap anywhere in Ecuador, placing them beyond the scope of in situ restoration. The geoSABINA geoportal extends this same logic to the cell scale, with the ranking directly available for any site in the territory.

Beyond species prioritization, the integration of climatic persistence with ecosystem services provides a basis for identifying multifunctional native species that can contribute to biodiversity and ecosystem resilience while, where appropriate, supporting local livelihoods. This broadens the potential application of the framework from conventional forest restoration to tree-based restoration pathways within productive landscapes, including agroforestry and silvopastoral systems.

The framework presented is transferable to other megadiverse tropical contexts with similar data and resource constraints, and its operational implementation through the geoSABINA geoportal demonstrates that it is possible to close the gap between scientific modelling and territorial decision-making without requiring advanced technical capacities. For Ecuador, this approach provides a timely basis for translating the evolving national restoration agenda into spatially explicit decisions on where to restore and which native species to use. By complementing area-based restoration priorities with climate-informed and multifunctional species portfolios, the framework can contribute to a transition from hectares restored towards climate-resilient, multifunctional and bioculturally embedded landscapes.

## Authors contribution statement

Conceptualization: RJHF, JMB, BT, RGM. Data curation: RJHF, JMB, BT. Formal analyses: RJHF, RGM, BT. Funding acquisition: RJHF, RGM. Supervision: RGM, BT. Writing – original draft: RJHF, JMB, RGM. Writing – review & editing: All authors.

## Acknowledgements

This work was partially supported by the CONNECT-Wild project (IAP-RP-AMR25005), an initiative of the International Academic Program (IAP) in collaboration with the David Rockefeller Center for Latin American Studies (DRCLAS), Harvard University, with institutional support from Universidad Autónoma de Madrid (UAM).

## Data availability

Species-level results are provided in Data S1 and the Services values used to compute the IPR in Data S2. Current and future climatic suitability and IPR values for each species and cell can be visualized and downloaded from the geoSABINA geoportal (https://www.geosabina.com). Occurrence records were obtained from GBIF, BIEN and SISBIO.

## REFERENCES

Adde, A., Külling, N., Rey, P., Fopp, F., Brun, P., Broennimann, O., Lehmann, A., Petitpierre, B., Zimmermann, N. E., Pellissier, L., Altermatt, F., & Guisan, A. (2024). Projecting Untruncated Climate Change Effects on Species’ Climate Suitability: Insights From an Alpine Country. Global Change Biology, 30(11), e17557. 10.1111/gcb.17557

Adde, A., Rey, P., Brun, P., Külling, N., Fopp, F., Altermatt, F., Broennimann, O., Lehmann, A., Petitpierre, B., Zimmermann, N. E., Pellissier, L., & Guisan, A. (2023). N-SDM: a high-performance computing pipeline for Nested Species Distribution Modelling. Ecography, 2023(6), e06540. 10.1111/ecog.06540

Adde, A., Rey, P.-L., Fopp, F., Petitpierre, B., Schweiger, A. K., Broennimann, O., Lehmann, A., Zimmermann, N. E., Altermatt, F., Pellissier, L., & Guisan, A. (2023). Too many candidates: Embedded covariate selection procedure for species distribution modelling with the covsel R package. Ecological Informatics, 75, 102080. 10.1016/j.ecoinf.2023.102080

Aguirre-Gutiérrez, J., Díaz, S., Rifai, S. W., Corral-Rivas, J. J., Nava-Miranda, M. G., González-M, R., Hurtado-M, A. B., Revilla, N. S., Vilanova, E., Almeida, E., De Oliveira, E. A., Alvarez-Davila, E., Alves, L. F., De Andrade, A. C. S., Lola Da Costa, A. C., Vieira, S. A., Aragão, L., Arets, E., Aymard C., G. A., … Malhi, Y. (2025). Tropical forests in the Americas are changing too slowly to track climate change. Science, 387(6738), eadl5414. 10.1126/science.adl5414

Allouche, O., Tsoar, A., & Kadmon, R. (2006). Assessing the accuracy of species distribution models: Prevalence, kappa and the true skill statistic (TSS). Journal of Applied Ecology, 43(6), 1223–1232. 10.1111/j.1365-2664.2006.01214.x

Andres, S. E., Atkinson, J., Coleman, D., Brazill-Boast, J., Wright, I. J., Allen, S., & Gallagher, R. V. (2024). Constraints of commercially available seed diversity in restoration: Implications for plant functional diversity. PLANTS, PEOPLE, PLANET, 6(6), 1341–1357. 10.1002/ppp3.10523

Anselmetto, N., Morresi, D., Barbarino, S., Loglisci, N., Betts, M. G., & Garbarino, M. (2025). Species distribution models built with local species data perform better for current time, but suffer from niche truncation. Agricultural and Forest Meteorology, 362, 110361. 10.1016/j.agrformet.2024.110361

Aragon-Osejo, J., Beltrán, L., Iglesias, J., Zambrano, H., Borja, D., Oñate, C., Simbaña, F., Valencia, F., Rodríguez De La Vera, K., Poveda, L., Proaño, F., Parra, L., Chacón, M., Llactayo, W., Pequeño Saco, T., Huamani, W., Marchand, G., Tinoco, R., Quispe, L., … Venter, O. (2025). National human footprint maps for Peru and Ecuador. Scientific Data, 13(1), 12. 10.1038/s41597-025-06301-0

Araujo, M., & New, M. (2007). Ensemble forecasting of species distributions. Trends in Ecology & Evolution, 22(1), 42–47. 10.1016/j.tree.2006.09.010

Balslev, H. (1988). DISTRIBUTION PATTERNS OF ECUADOREAN PLANT SPECIES. TAXON, 37(3), 567–577. 10.2307/1221100

Banks, K. M., Edwards, O. M., Zhang, B., & Reichert, M. S. (2026). Using spatially-nested hierarchical species distribution models to estimate current and future distributions of a cryptic species at a regional scale. Journal of Animal Ecology, 95(5), 747–762. 10.1111/1365-2656.70248

Barbet-Massin, M., Jiguet, F., Albert, C. H., & Thuiller, W. (2012). Selecting pseudo-absences for species distribution models: How, where and how many? Methods in Ecology and Evolution, 3(2), 327–338. 10.1111/j.2041-210X.2011.00172.x

Barragán, G., & Rhemtulla, J. M. (2025). Local people in both frontier and intensifying landscapes in Ecuador desire forest restoration that provides multiple benefits. People and Nature, 7(8), 1922–1939. 10.1002/pan3.70084

Barragán, G., Wang, T., & Rhemtulla, J. M. (2023). Trees planted under a global restoration pledge have mixed futures under climate change. Restoration Ecology, 31(3), e13764. 10.1111/rec.13764

Bass, M. S., Finer, M., Jenkins, C. N., Kreft, H., Cisneros-Heredia, D. F., McCracken, S. F., Pitman, N. C. A., English, P. H., Swing, K., Villa, G., Di Fiore, A., Voigt, C. C., & Kunz, T. H. (2010). Global Conservation Significance of Ecuador’s Yasuní National Park. PLoS ONE, 5(1), e8767. 10.1371/journal.pone.0008767

Booth, T. H. (2022). Checking bioclimatic variables that combine temperature and precipitation data before their use in species distribution models. Austral Ecology, 47(7), 1506–1514. 10.1111/aec.13234

Brancalion, P. H. S., & Holl, K. D. (2020). Guidance for successful tree planting initiatives. Journal of Applied Ecology, 57(12), 2349–2361. 10.1111/1365-2664.13725

Brancalion, P. H. S., Hua, F., Joyce, F. H., Antonelli, A., & Holl, K. D. (2025). Moving biodiversity from an afterthought to a key outcome of forest restoration. Nature Reviews Biodiversity, 1(4), 248–261. 10.1038/s44358-025-00032-1

Breiman, L. (2001). Random Forests. Machine Learning, 45(1), 5–32. 10.1023/A:1010933404324

Brodie, S., Smith, J. A., Muhling, B. A., Barnett, L. A. K., Carroll, G., Fiedler, P., Bograd, S. J., Hazen, E. L., Jacox, M. G., Andrews, K. S., Barnes, C. L., Crozier, L. G., Fiechter, J., Fredston, A., Haltuch, M. A., Harvey, C. J., Holmes, E., Karp, M. A., Liu, O. R., … Kaplan, I. C. (2022). Recommendations for quantifying and reducing uncertainty in climate projections of species distributions. Global Change Biology, 28(22), 6586–6601. 10.1111/gcb.16371

Broennimann, O., & Guisan, A. (2026). High-resolution topoclimate data is essential for predicting species distributions in mountains: Insights from CHclim25. Alpine Botany. 10.1007/s00035-026-00365-3

Carrillo-García, C., Girola-Iglesias, L., Guijarro, M., Hernando, C., Madrigal, J., & Mateo, R. G. (2023). Ecological niche models applied to post-megafire vegetation restoration in the context of climate change. Science of The Total Environment, 855, 158858. 10.1016/j.scitotenv.2022.158858

Chacón-Moreno, E., Alía, R., Fernández-Pastor, M., Gaitán, E., García Del Barrio, J. M., Mateo, R. G., García Pose, A., Grivet, D., Guadaño-Peyrot, C., Hernández-Parada, A., Morales-Barbero, J., Paradinas Blázquez, C., Prado López, C., Rey-Benayas, J. M., Ribalaygua, J., Rodríguez-Ruiz, G., Royé, D., Olsson, S., & Del Carre Díaz, M. (2026). A framework for climate-resilient forest planning and restoration: Advances linking species distribution modelling, genetic adaptation and future climate scenarios. Frontiers in Forests and Global Change, 9, 1735354. 10.3389/ffgc.2026.1735354

Chave, J., Coomes, D., Jansen, S., Lewis, S. L., Swenson, N. G., & Zanne, A. E. (2009). Towards a worldwide wood economics spectrum. Ecology Letters, 12(4), 351–366. 10.1111/j.1461-0248.2009.01285.x

Chevalier, M., Broennimann, O., Cornuault, J., & Guisan, A. (2021). Data integration methods to account for spatial niche truncation effects in regional projections of species distribution. Ecological Applications, 31(7), e02427. 10.1002/eap.2427

Chevalier, M., Zarzo-Arias, A., Guélat, J., Mateo, R. G., & Guisan, A. (2022). Accounting for niche truncation to improve spatial and temporal predictions of species distributions. Frontiers in Ecology and Evolution, 10, 944116. 10.3389/fevo.2022.944116

Cogălniceanu, D., & Szekely, D. (2025). The hidden costs of gold mining in Ecuador. Science, 388(6752), 1143–1143. 10.1126/science.adx3004

Collart, F., & Guisan, A. (2023). Small to train, small to test: Dealing with low sample size in model evaluation. Ecological Informatics, 75, 102106. 10.1016/j.ecoinf.2023.102106

Daru, B. H. (2025). Tracking hidden dimensions of plant biogeography from herbaria. New Phytologist, 246(1), 61–77. 10.1111/nph.70002

De Freitas, P. D., & Junior, P. M. G. (2025). Why Study Biodiversity in the Neotropics? En M. V. Bohrer Monteiro Siqueira, E. R. Konzen, & P. M. Galetti Junior (Eds.), Population Genetics in the Neotropics (pp. 19–40). Springer Nature Switzerland. 10.1007/978-3-031-83685-5_2

de la Torre, Lucía, Navarrete, Hugo, Muriel M., Priscilla, Macía Barco, Manuel Juan, & Balslev, Henrik (Eds.). (2008). Enciclopedia de las plantas útiles del Ecuador (1. ed). Herbario QCA de la Escuela de Ciencias Biológicas de la Pontificia Universidad Católica del Ecuador ; Herbario AAU del Departamento de Ciencias Biológicas de la Universidad de Aarhus.

Di Sacco, A., Hardwick, K. A., Blakesley, D., Brancalion, P. H. S., Breman, E., Cecilio Rebola, L., Chomba, S., Dixon, K., Elliott, S., Ruyonga, G., Shaw, K., Smith, P., Smith, R. J., & Antonelli, A. (2021). Ten golden rules for reforestation to optimize carbon sequestration, biodiversity recovery and livelihood benefits. Global Change Biology, 27(7), 1328–1348. 10.1111/gcb.15498

Dogan, H., & Yilmaz, O. Y. (2026). Nested species distribution models highlight scale and climate-dataset effects on adaptation planning for sweet chestnut (Castanea sativa) forests in Türkiye. In Review. 10.21203/rs.3.rs-8708047/v1

Edwards, D. P., & Cerullo, G. R. (2024). Biodiversity is central for restoration. Current Biology, 34(9), R371–R379. 10.1016/j.cub.2024.02.032

Engemann, K., Enquist, B. J., Sandel, B., Boyle, B., Jørgensen, P. M., Morueta-Holme, N., Peet, R. K., Violle, C., & Svenning, J. (2015). Limited sampling hampers “big data” estimation of species richness in a tropical biodiversity hotspot. Ecology and Evolution, 5(3), 807–820. 10.1002/ece3.1405

Enquist, B. J., Boyle, B., Maitner, B. S., Merow, C., Carlson, B. S., Feng, X., Moulatlet, G. M., Newman, E. A., Park, D. S., Pillet, M., Roehrdanz, P. R., Serra-Diaz, J. M., Babu, R. S. Y., Barbosa, G. G. C., Casler, N., Donoghue, J. C., Guaderrama, D., Sajja, R. K., Couvreur, T. L. P., … Wiser, S. K. (2026). BIEN: A biodiversity informatics ecosystem advancing open and reproducible workflows for plant observation, plot and trait data. Methods in Ecology and Evolution, 17(5), 1556–1584. 10.1111/2041-210x.70274

Enquist, B. J., Feng, X., Boyle, B., Maitner, B., Newman, E. A., Jørgensen, P. M., Roehrdanz, P. R., Thiers, B. M., Burger, J. R., Corlett, R. T., Couvreur, T. L. P., Dauby, G., Donoghue, J. C., Foden, W., Lovett, J. C., Marquet, P. A., Merow, C., Midgley, G., Morueta-Holme, N., … McGill, B. J. (2019). The commonness of rarity: Global and future distribution of rarity across land plants. Science Advances, 5(11), eaaz0414. 10.1126/sciadv.aaz0414

Erbaugh, J. T., Pradhan, N., Adams, J., Oldekop, J. A., Agrawal, A., Brockington, D., Pritchard, R., & Chhatre, A. (2020). Global forest restoration and the importance of prioritizing local communities. Nature Ecology & Evolution, 4(11), 1472–1476. 10.1038/s41559-020-01282-2

Erickson, V. J., & Halford, A. (2020). Seed planning, sourcing, and procurement. Restoration Ecology, 28(S3). 10.1111/rec.13199

Fadrique, B., Costa, F., Cuesta, F., Arellano, G., Cayuela, L., Baker, T. R., Draper, F. C., Esquivel-Muelbert, A., Ter Steege, H., Bauters, M., Aguirre-Gutiérrez, J., Aguirre-Mendoza, Z., Alexiades, M. N., Alvarez-Davila, E., Arets, E., Ayala, E., Aymard, C. G. A., Baccaro, F., Báez, S., … Phillips, O. L. (2026). Tree diversity is changing across tropical Andean and Amazonian forests in response to global change. Nature Ecology & Evolution, 10(2), 267–280. 10.1038/s41559-025-02956-5

Fajardo, J., Lessmann, J., Devenish, C., Bonaccorso, E., Felicísimo, Á. M., Rojas-Runjaic, F. J. M., Rojas, H., Lentino, M., Muñoz, J., & Mateo, R. G. (2023). The performance of protected-area expansions in representing tropical Andean species: Past trends and climate change prospects. Scientific Reports, 13(1), 966. 10.1038/s41598-022-27365-7

Farfan-Rios, W., Feeley, K. J., Myers, J. A., Tello, S., Sallo-Bravo, J., Malhi, Y., Phillips, O. L., Baker, T. R., Nina-Quispe, A., Garcia-Cabrera, K., Saatchi, S. S., Terborgh, J. W., Pitman, N. C. A., Monteagudo Mendoza, A. L., Vasquez, R., Salinas, N., Cayola, L., Fuentes Claros, A., Loza, M. I., … Silman, M. R. (2025). Amazonian and Andean tree communities are not tracking current climate warming. Proceedings of the National Academy of Sciences, 122(34), e2425619122. 10.1073/pnas.2425619122

FBA. (2026). Fondo Biocorredor Amazónico. Fondo Biocorredor Amazónico. https://fondobiocorredoramazonico.org/biocorredor-amazonico/

Feeley, K. J., Bernal-Escobar, M., Fortier, R., & Kullberg, A. T. (2023). Tropical Trees Will Need to Acclimate to Rising Temperatures—But Can They? Plants, 12(17), 3142. 10.3390/plants12173142

Fick, S. E., & Hijmans, R. J. (2017). WorldClim 2: New 1-km spatial resolution climate surfaces for global land areas. International Journal of Climatology, 37(12), 4302–4315. 10.1002/joc.5086

Fielding, A. H., & Bell, J. F. (1997). A review of methods for the assessment of prediction errors in conservation presence/absence models. Environmental Conservation, 24(1), 38–49. 10.1017/S0376892997000088

Franklin, J. (2023). Species distribution modelling supports the study of past, present and future biogeographies. Journal of Biogeography, 50(9), 1533–1545. 10.1111/jbi.14617

Freire-Fierro, A., Tirira, D., Batallas, R., Oleas, N., Rodríguez, C., Vogt, C., Cadena-Ortiz, H., Ulloa Ulloa, C., Salazar, F., & Donoso, D. (2024). Ecuadorian Red Lists and the IUCN Red List: A Disconnect. Biodiversity Information Science and Standards, 8, e138924. 10.3897/biss.8.138924

Fremout, T., Thomas, E., Gaisberger, H., Van Meerbeek, K., Muenchow, J., Briers, S., Gutierrez-Miranda, C. E., Marcelo-Peña, J. L., Kindt, R., Atkinson, R., Cabrera, O., Espinosa, C. I., Aguirre-Mendoza, Z., & Muys, B. (2020). Mapping tree species vulnerability to multiple threats as a guide to restoration and conservation of tropical dry forests. Global Change Biology, 26(6), 3552–3568. 10.1111/gcb.15028

Fremout, T., Thomas, E., Taedoumg, H., Briers, S., Gutiérrez-Miranda, C. E., Alcázar-Caicedo, C., Lindau, A., Mounmemi Kpoumie, H., Vinceti, B., Kettle, C., Ekué, M., Atkinson, R., Jalonen, R., Gaisberger, H., Elliott, S., Brechbühler, E., Ceccarelli, V., Krishnan, S., Vacik, H., … Muys, B. (2022). Diversity for Restoration (D4R): Guiding the selection of tree species and seed sources for climate-resilient restoration of tropical forest landscapes. Journal of Applied Ecology, 59(3), 664–679. 10.1111/1365-2664.14079

Galindo, V., Calle, Z., Chará, J., & Armbrecht, I. (2017). Facilitation by pioneer shrubs for the ecological restoration of riparian forests in the Central Andes of Colombia. Restoration Ecology, 25(5), 731–737. 10.1111/rec.12490

García Del Barrio, J. M., Guadaño-Peyrot, C., Grivet, D., Olsson, S., Barba, D., Auñon, F., Chacón-Moreno, E., Fernández-Pastor, M., Gaitán, E., Mateo, R. G., García-Pose, A., Hernández-Parada, A., Morales-Barbero, J., Paradinas-Blázquez, C., Prado-López, C., Rey-Benayas, J. M., Ribalaygua, J., Rodríguez-Ruiz, G., Royé, D., … Alía, R. (2026). Vulnerability to Climate Change of Forest Genetic Conservation Units Assessed Through Species Distribution Models. Land, 15(5), 890. 10.3390/land15050890

Gastón, A., García-Viñas, J. I., Bravo-Fernández, A. J., López-Leiva, C., Oliet, J. A., Roig, S., & Serrada, R. (2014). Species distribution models applied to plant species selection in forest restoration: Are model predictions comparable to expert opinion? New Forests, 45(5), 641–653. 10.1007/s11056-014-9427-7

Gaul, W., Sadykova, D., White, H. J., Leon-Sanchez, L., Caplat, P., Emmerson, M. C., & Yearsley, J. M. (2020). Data quantity is more important than its spatial bias for predictive species distribution modelling. PeerJ, 8, e10411. 10.7717/peerj.10411

Gittleman, J. L., Pimm, S. L., Van Rensburg, B. J., Van Houtan, K. S., & Roberts, C. M. (2026). Implementing the Kunming-Montreal Biodiversity Targets. Proceedings of the National Academy of Sciences, 123(2), e2514761123. 10.1073/pnas.2514761123

Goicolea, T., Adde, A., Broennimann, O., García-Viñas, J. I., Gastón, A., José Aroca-Fernández, M., Guisan, A., & G. Mateo, R. (2025). Spatially-nested hierarchical species distribution models to overcome niche truncation in national-scale studies. Ecography, 2025(3), e07328. 10.1111/ecog.07328

Goicolea, T., Morales-Barbero, J., García-Viñas, J. I., Gastón, A., Aroca-Fernández, M. J., Calleja, J. A., Moreno, J. C., Ramos-Gutiérrez, I., Rodríguez, M. Á., Lima, H., Broennimann, O., Guisan, A., Adde, A., Pérez-Latorre, A. V., & Mateo, R. G. (2026). A unified plant ecology database for Spain. Scientific Data, 13(1), 355. 10.1038/s41597-026-06757-8

Guevara, E. A., Satama-Bermeo, M., Quinteros, L., Santander, T., Olschewski, R., & Farwig, N. (2026). Understanding landowners’ preferences to participate in tropical mountain forest restoration. Restoration Ecology, 34(5), e70368. 10.1111/rec.70368

Guisan, A., Chevalier, M., Adde, A., Zarzo-Arias, A., Goicolea, T., Broennimann, O., Petitpierre, B., Scherrer, D., Rey, P., Collart, F., Riva, F., Steen, B., & Mateo, R. G. (2025). Spatially nested species distribution models (N-SDM): An effective tool to overcome niche truncation for more robust inference and projections. Journal of Ecology, 113(7), 1588–1605. 10.1111/1365-2745.70063

Guisan, A., Thuiller, W., & Zimmermann, N. E. (with Di Cola, V., Georges, D., & Psomas, A.). (2017). Habitat suitability and distribution models: With applications in R. Cambridge University Press.

Guzmán, N., Gentili, R., Jiménez, M., Ansaloni, R., Citterio, S., & Minga, D. (2026). Exploring plant α- and β-diversity patterns along tropical altitudinal gradients in the Southern Andes of Ecuador: A potential pathway to biodiversity conservation. Plant Biosystems, 160(3), 159. 10.1007/s44473-026-00165-x

Hastie, T. J., & Hastie, T. J. (1991). Statistical Models in S (1st ed). CRC Press.

Hellegers, M., van Hinsberg, A., Lenoir, J., Dengler, J., Huijbregts, M. A. J., & Schipper, A. M. (2025). Multiple Threshold-Selection Methods Are Needed to Binarise Species Distribution Model Predictions. Diversity and Distributions, 31(4), e70019. 10.1111/ddi.70019

Hernandez-Montilla, M., Devenish, K., Alencar, L., Benzeev, R., Choksi, P., Djenontin, I. N. S., Fagan, M. E., Fischer, H., Isaacs Cubides, P., Linhares Juvenal, T., Kabutey-Ongor, M., Kamoto, J., Kinzer, A., Kroeker-Maus, D., Mansourian, S., Melo, F., Miller, D. C., Nofyanza, S., Pradeep, A., … Oldekop, J. A. (2026). Forwarding forest restoration: Seven key socio-ecological issues for advancing forest restoration in a world in flux. People and Nature, 8(3), 569–581. 10.1002/pan3.70254

Herrera-Feijoo, R. J., Torres, B., López-Tobar, R., Tipán-Torres, C., Toulkeridis, T., Heredia-R, M., & Mateo, R. G. (2023). Modelling Climatically Suitable Areas for Mahogany (Swietenia macrophylla King) and Their Shifts across Neotropics: The Role of Protected Areas. Forests, 14(2), 385. 10.3390/f14020385

Hirzel, A. H., Le Lay, G., Helfer, V., Randin, C., & Guisan, A. (2006). Evaluating the ability of habitat suitability models to predict species presences. Ecological Modelling, 199(2), 142–152. 10.1016/j.ecolmodel.2006.05.017

Holl, K. D., Luong, J. C., & Brancalion, P. H. S. (2022). Overcoming biotic homogenization in ecological restoration. Trends in Ecology & Evolution, 37(9), 777–788. 10.1016/j.tree.2022.05.002

Hughes, A., Demirel, N., Barnes, D., Ahlquist, I., Ondo, I., Guralnick, R., Hirsch, T., Enquist, B., Merow, C., Kaschner, K., Reygondeau, G., Egorova, Y., Orr, M., Qiao, H., Stephenson, P., Waller, J., & Burgess, N. (2025). Understanding the adequacy and representativeness of species distribution data. Ecology and Evolutionary Biology. 10.32942/X28S82

Humphreys, A. M., Fisher, D. O., Witts, N. A., Silvestro, D., & Antonelli, A. (2025). Harnessing the benefits of herbarium specimen digitisation for inferring recent and ongoing plant extinctions. New Phytologist, nph.70552. 10.1111/nph.70552

Hutchinson, A., Zito, A. R., & McGowan, P. J. K. (2026). Pathways for transforming biodiversity governance: An examination of the Global Biodiversity Framework’s Considerations. Ambio, 55(1), 56–67. 10.1007/s13280-025-02215-8

Jensen, D. A., Rao, M., Zhang, J., Grøn, M., Tian, S., Ma, K., & Svenning, J.-C. (2021). The potential for using rare, native species in reforestation– A case study of yews (Taxaceae) in China. Forest Ecology and Management, 482, 118816. 10.1016/j.foreco.2020.118816

Kass, J. M., Smith, A. B., Warren, D. L., Vignali, S., Schmitt, S., Aiello-Lammens, M. E., Arlé, E., Márcia Barbosa, A., Broennimann, O., Cobos, M. E., Guéguen, M., Guisan, A., Merow, C., Naimi, B., Nobis, M. P., Ondo, I., Osorio-Olvera, L., Owens, H. L., Pinilla-Buitrago, G. E., … Zurell, D. (2025). Achieving higher standards in species distribution modeling by leveraging the diversity of available software. Ecography, 2025(2), e07346. 10.1111/ecog.07346

Ke, G., Meng, Q., Finley, T., Wang, T., Chen, W., Ma, W., Ye, Q., & Liu, T.-Y. (2017). LightGBM: A Highly Efficient Gradient Boosting Decision Tree. En I. Guyon, U. V. Luxburg, S. Bengio, H. Wallach, R. Fergus, S. Vishwanathan, & R. Garnett (Eds.), Advances in Neural Information Processing Systems (Vol. 30). Curran Associates, Inc. https://proceedings.neurips.cc/paper_files/paper/2017/file/6449f44a102fde848669bdd9eb6b76fa-Paper.pdf

Krainovic, P. M., Romanelli, J. P., De Resende, A. F., Simões, L. H. P., Boeni, A. F., Souza, L. R., Toledo, C. A. P., Souza, V. C., Guillemot, J., Brouwer, R., Bongers, F., Molin, P. G., Chaves, R. B., Massi, K. G., Rodrigues, R. R., de-Miguel, S., & Brancalion, P. H. S. (2025). Bioeconomic opportunities in restored tropical forests. Ambio. 10.1007/s13280-025-02234-5

Lacoretz, M. V., & Pérez, D. R. (2025). Integrating theory and practice: Knowledge exchange and engagement of nursery managers in ecological restoration. Restoration Ecology, 33(1), e14282. 10.1111/rec.14282

Lessmann, J., Muñoz, J., & Bonaccorso, E. (2014). Maximizing species conservation in continental E cuador: A case of systematic conservation planning for biodiverse regions. Ecology and Evolution, 4(12), 2410–2422. 10.1002/ece3.1102

Li, G., Fang, C., Watson, J. E. M., Sun, S., Qi, W., Wang, Z., & Liu, J. (2024). Mixed effectiveness of global protected areas in resisting habitat loss. Nature Communications, 15(1), 8389. 10.1038/s41467-024-52693-9

Lobos, R. B. C., Ricci, L., Martini, M., Testolin, R., Di Musciano, M., Moudrý, V., Groom, Q., & Rocchini, D. (2026). When accessibility distorts niche estimates: A virtual species framework across species prevalence and sample size gradients. Ecological Informatics, 97, 103937. 10.1016/j.ecoinf.2026.103937

López-Tobar, R., García Robredo, F., & Torres Navarrete, B. (2026). Tipología de especies maderables comerciales y potencial económico de su aprovechamiento en la Amazonía ecuatoriana. Colombia forestal, 29(1), e23680. 10.14483/2256201X.23680

López-Tobar, R., Herrera-Feijoo, R. J., García-Robredo, F., Mateo, R. G., & Torres, B. (2024). Timber harvesting and conservation status of forest species in the Ecuadorian Amazon. Frontiers in Forests and Global Change, 7, 1389852. 10.3389/ffgc.2024.1389852

López-Tobar, R., Herrera-Feijoo, R. J., Mateo, R. G., García-Robredo, F., & Torres, B. (2023). Botanical Collection Patterns and Conservation Categories of the Most Traded Timber Species from the Ecuadorian Amazon: The Role of Protected Areas. Plants, 12(18), 3327. 10.3390/plants12183327

Lortie, C. J., Filazzola, A., Westphal, M., & Butterfield, H. S. (2022). Foundation plant species provide resilience and microclimatic heterogeneity in drylands. Scientific Reports, 12(1), 18005. 10.1038/s41598-022-22579-1

MAE. (2019). Plan Nacional deRestauración Forestal 2019—2030. Ministerio de Ambiente y Energía del Ecuador. https://initiative20x20.org/sites/default/files/2021-08/MAE_2019_Plan%20Nacional_Restauracion%20%282%29%20%281%29.pdf

MAE. (2023). Deforestación y regeneración a nivel provincial del periodo 1990-2022 del Ecuador continental. Infraestructura de Datos Especial del MAATE. https://ide.ambiente.gob.ec/mapainteractivo/

Malhotra, A., Von Fromm, S. F., Bond-Lamberty, B., Doetterl, S., Georgiou, K., Graham, E. B., Heckman, K. A., Jia, R., Patel, K. F., Rod, K. A., Santos, F., Terrer, C., Todd-Brown, K., Zheng, J., Hofmockel, K., & Bailey, V. (2026). Do we have globally representative data to understand soil processes? Biogeochemistry, 169(2), 23. 10.1007/s10533-025-01301-z

Malizia, A., Blundo, C., Carilla, J., Osinaga Acosta, O., Cuesta, F., Duque, A., Aguirre, N., Aguirre, Z., Ataroff, M., Baez, S., Calderón-Loor, M., Cayola, L., Cayuela, L., Ceballos, S., Cedillo, H., Farfán Ríos, W., Feeley, K. J., Fuentes, A. F., Gámez Álvarez, L. E., … Young, K. R. (2020). Elevation and latitude drives structure and tree species composition in Andean forests: Results from a large-scale plot network. PLOS ONE, 15(4), e0231553. 10.1371/journal.pone.0231553

Marmion, M., Parviainen, M., Luoto, M., Heikkinen, R. K., & Thuiller, W. (2009). Evaluation of consensus methods in predictive species distribution modelling. Diversity and Distributions, 15(1), 59–69. 10.1111/j.1472-4642.2008.00491.x

Marshall, A. R., Waite, C. E., Pfeifer, M., Banin, L. F., Rakotonarivo, S., Chomba, S., Herbohn, J., Gilmour, D. A., Brown, M., & Chazdon, R. L. (2023). Fifteen essential science advances needed for effective restoration of the world’s forest landscapes. Philosophical Transactions of the Royal Society B: Biological Sciences, 378(1867), 20210065. 10.1098/rstb.2021.0065

Martin, M. P., Woodbury, D. J., Doroski, D. A., Nagele, E., Storace, M., Cook-Patton, S. C., Pasternack, R., & Ashton, M. S. (2021). People plant trees for utility more often than for biodiversity or carbon. Biological Conservation, 261, 109224. 10.1016/j.biocon.2021.109224

Mateo, R. G., De La Estrella, M., Felicísimo, Á. M., Muñoz, J., & Guisan, A. (2013). A new spin on a compositionalist predictive modelling framework for conservation planning: A tropical case study in Ecuador. Biological Conservation, 160, 150–161. 10.1016/j.biocon.2013.01.014

Mateo, R. G., Felicísimo, Á. M., & Muñoz, J. (2010). Effects of the number of presences on reliability and stability of MARS species distribution models: The importance of regional niche variation and ecological heterogeneity: Effects of the number of presences on model reliability and stability. Journal of Vegetation Science, 21(5), 908–922. 10.1111/j.1654-1103.2010.01198.x

Mateo, R. G., Felicísimo, Á. M., Pottier, J., Guisan, A., & Muñoz, J. (2012). Do Stacked Species Distribution Models Reflect Altitudinal Diversity Patterns? PLoS ONE, 7(3), e32586. 10.1371/journal.pone.0032586

Mateo, R. G., Gastón, A., Aroca-Fernández, M. J., Broennimann, O., Guisan, A., Saura, S., & García-Viñas, J. I. (2019). Hierarchical species distribution models in support of vegetation conservation at the landscape scale. Journal of Vegetation Science, 30(2), 386–396. 10.1111/jvs.12726

Mateo, R. G., Morales-Barbero, J., Zarzo-Arias, A., Lima, H., Gómez-Rubio, V., & Goicolea, T. (2024). sabinaNSDM: An R package for spatially nested hierarchical species distribution modelling. Methods in Ecology and Evolution, 15(10), 1796–1803. 10.1111/2041-210X.14417

Meinshausen, M., Nicholls, Z. R. J., Lewis, J., Gidden, M. J., Vogel, E., Freund, M., Beyerle, U., Gessner, C., Nauels, A., Bauer, N., Canadell, J. G., Daniel, J. S., John, A., Krummel, P. B., Luderer, G., Meinshausen, N., Montzka, S. A., Rayner, P. J., Reimann, S., … Wang, R. H. J. (2020). The shared socio-economic pathway (SSP) greenhouse gas concentrations and their extensions to 2500. Geoscientific Model Development, 13(8), 3571–3605. 10.5194/gmd-13-3571-2020

Mestre-Tomás, J. (2025). GeoThinneR: An R Package for Efficient Spatial Thinning of Species Occurrences and Point Data (arXiv:2505.07867). arXiv. 10.48550/arXiv.2505.07867

Ministerio de Ambiente y Energía (MAE), Ministerio de Agricultura, Ganadería y Pesca (MAGP), & Ministerio de. (2026). Estrategia Forestal Sostenible del Ecuador (EFSE) 2026-2050. Gobierno del Ecuador.

Moreno-Amat, E., Mateo, R. G., Nieto-Lugilde, D., Morueta-Holme, N., Svenning, J.-C., & García-Amorena, I. (2015). Impact of model complexity on cross-temporal transferability in Maxent species distribution models: An assessment using paleobotanical data. Ecological Modelling, 312, 308–317. 10.1016/j.ecolmodel.2015.05.035

Moudrý, V., Bazzichetto, M., Remelgado, R., Devillers, R., Lenoir, J., Mateo, R. G., Lembrechts, J. J., Sillero, N., Lecours, V., Cord, A. F., Barták, V., Balej, P., Rocchini, D., Torresani, M., Arenas-Castro, S., Man, M., Prajzlerová, D., Gdulová, K., Prošek, J., … Šímová, P. (2024). Optimising occurrence data in species distribution models: Sample size, positional uncertainty, and sampling bias matter. Ecography, 2024(12), e07294. 10.1111/ecog.07294

Mouillot, D., Bellwood, D. R., Baraloto, C., Chave, J., Galzin, R., Harmelin-Vivien, M., Kulbicki, M., Lavergne, S., Lavorel, S., Mouquet, N., Paine, C. E. T., Renaud, J., & Thuiller, W. (2013). Rare Species Support Vulnerable Functions in High-Diversity Ecosystems. PLoS Biology, 11(5), e1001569. 10.1371/journal.pbio.1001569

Neeson, T. M., Doran, P. J., Ferris, M. C., Fitzpatrick, K. B., Herbert, M., Khoury, M., Moody, A. T., Ross, J., Yacobson, E., & McIntyre, P. B. (2018). Conserving rare species can have high opportunity costs for common species. Global Change Biology, 24(8), 3862–3872. 10.1111/gcb.14162

Neill, D. A. (2012). ¿Cuántas especies nativas de plantas vasculares hay en Ecuador? Revista Amazónica. Ciencia y Tecnología, 1(1), 70–83. 10.59410/RACYT-v01n01ep08-0001

Owusu Danquah, E., Frimpong, F., Boadi, S., Peprah, C. O., Aduhene-Chinbuah, J., Opoku, P., Danquah, F. O., Baafi, E., Botey, H. M., Aidoo, K. A. S., Acheampong, P. P., Pokuaa, Y., & Asante, M. D. (2026). Forest and biodiversity conservation for food and livelihood security: A review of opportunities and challenges of community-based approaches. Frontiers in Forests and Global Change, 9, 1890643. 10.3389/ffgc.2026.1890643

Poggio, L., De Sousa, L. M., Batjes, N. H., Heuvelink, G. B. M., Kempen, B., Ribeiro, E., & Rossiter, D. (2021). SoilGrids 2.0: Producing soil information for the globe with quantified spatial uncertainty. SOIL, 7(1), 217–240. 10.5194/soil-7-217-2021

Porfirio, L. L., Harris, R. M. B., Lefroy, E. C., Hugh, S., Gould, S. F., Lee, G., Bindoff, N. L., & Mackey, B. (2014). Improving the Use of Species Distribution Models in Conservation Planning and Management under Climate Change. PLoS ONE, 9(11), e113749. 10.1371/journal.pone.0113749

Qiao, H., & Escobar, L. E. (2025). A practical framework for a theory-driven ecological niche modeling workflow. The Innovation Life, 3(4), 100165. 10.59717/j.xinn-life.2025.100165

Raven, P. H., Gereau, R. E., Phillipson, P. B., Chatelain, C., Jenkins, C. N., & Ulloa Ulloa, C. (2020). The distribution of biodiversity richness in the tropics. Science Advances, 6(37), eabc6228. 10.1126/sciadv.abc6228

Réjou-Méchain, M., Ariane Tanguy, Piponiot, C., Chave, J., & Hérault, B. (2017). biomass: An r package for estimating above-ground biomass and its uncertainty in tropical forests. Methods in Ecology and Evolution, 8(9), 1163–1167. 10.1111/2041-210X.12753

Restrepo, Z., González-Caro, S., Hartley, I. P., Villegas, J. C., Meir, P., Sanchez, A., Ruiz Carrascal, D., & Mercado, L. M. (2025). Heterogeneous thermal tolerance of dominant Andean montane tree species. Communications Earth & Environment, 6(1), 117. 10.1038/s43247-025-02083-w

Richter, M., Diertl, K.-H., Emck, P., Peters, T., & Beck, E. (2009). Reasons for an outstanding plant diversity in the tropical Andes of Southern Ecuador. Landscape Online, 12, 1–35. 10.3097/LO.200912

Rosenfield, M. F., Jardim, L., Antongiovanni, M., Querido, L. C. D. A., Ribeiro, A. A., Sánchez-Tapia, A., Silveira, P., Terribile, L. C., Venticinque, E. M., Albernaz, A. L., Garcia, L. C., Tambosi, L. R., Adami, M., Becker, F. G., Benchimol, M., Carvalheiro, L. G., Cornelius, C., Damasceno-Junior, G. A., Dobrovolski, R., … Garcia, E. (2025). Mapping Resilient Landscapes to Climate Change in a Megadiverse Country. Global Change Biology, 31(10), e70544. 10.1111/gcb.70544

Sanguet, A., Wyler, N., Petitpierre, B., Honeck, E., Poussin, C., Martin, P., & Lehmann, A. (2022). Beyond topo-climatic predictors: Does habitats distribution and remote sensing information improve predictions of species distribution models? Global Ecology and Conservation, 39, e02286. 10.1016/j.gecco.2022.e02286

Scherrer, D., D’Amen, M., Fernandes, R. F., Mateo, R. G., & Guisan, A. (2018). How to best threshold and validate stacked species assemblages? Community optimisation might hold the answer. Methods in Ecology and Evolution, 9(10), 2155–2166. 10.1111/2041-210X.13041

Song, L., Frazier, A. E., Roehrdanz, P. R., Yang, W., Krieger, J. M., McManus, N., Urbano, E. A. N., Buritica, S. R., & Willett, D. (2026). Modeling climate adaptive habitat connectivity for conservation planning in Colombia. Ecography, e08789. 10.1002/ecog.08789

Tebaldi, C., Debeire, K., Eyring, V., Fischer, E., Fyfe, J., Friedlingstein, P., Knutti, R., Lowe, J., O’Neill, B., Sanderson, B., Van Vuuren, D., Riahi, K., Meinshausen, M., Nicholls, Z., Tokarska, K. B., Hurtt, G., Kriegler, E., Lamarque, J.-F., Meehl, G., … Ziehn, T. (2021). Climate model projections from the Scenario Model Intercomparison Project (ScenarioMIP) of CMIP6. Earth System Dynamics, 12(1), 253–293. 10.5194/esd-12-253-2021

Thirisha Rani, D., Yagna Priya, M., Priyadharshini, S., & Devi, P. B. (2026). Conservation of Endemic Plants: Conventional and Biotechnological Approaches. En N. Kumar (Ed.), Biotechnology in Plant Conservation (pp. 25–43). Springer Nature Switzerland. 10.1007/978-3-032-02605-7_3

Thomas, E., Jalonen, R., Loo, J., Boshier, D., Gallo, L., Cavers, S., Bordács, S., Smith, P., & Bozzano, M. (2014). Genetic considerations in ecosystem restoration using native tree species. Forest Ecology and Management, 333, 66–75. 10.1016/j.foreco.2014.07.015

Thuiller, W. (2024). Ecological niche modelling. Current Biology, 34(6), R225–R229. 10.1016/j.cub.2024.02.018

Thuiller, W., Lafourcade, B., Engler, R., & Araújo, M. B. (2009). BIOMOD – a platform for ensemble forecasting of species distributions. Ecography, 32(3), 369–373. 10.1111/j.1600-0587.2008.05742.x

Torres, B., Espinoza, Í., Torres, A., Herrera-Feijoo, R., Luna, M., & García, A. (2023). Livelihood Capitals and Opportunity Cost for Grazing Areas’ Restoration: A Sustainable Intensification Strategy in the Ecuadorian Amazon. Animals, 13(4), 714. 10.3390/ani13040714

Torres, B., Tipán-Torres, C., Reyes, H., Tapia, A., Muñoz-Rengifo, J., Herrera-Feijoo, R., & García, A. (2026). Biocultural Productive Landscapes in the Andean–Amazon: Carbon, Biodiversity, and Livelihoods in Market-Linked Traditional Systems. Sustainability, 18(5), 2451. 10.3390/su18052451

Velásquez-Tibatá, J., Olaya-Rodríguez, M. H., López-Lozano, D., Gutiérrez, C., González, I., & Londoño-Murcia, M. C. (2019). BioModelos: A collaborative online system to map species distributions. PLOS ONE, 14(3), e0214522. 10.1371/journal.pone.0214522

Watson, J. E. M., Venegas-Li, R., Grantham, H., Dudley, N., Stolton, S., Rao, M., Woodley, S., Hockings, M., Burkart, K., Simmonds, J. S., Sonter, L. J., Sreekar, R., Possingham, H. P., & Ward, M. (2023). Priorities for protected area expansion so nations can meet their Kunming-Montreal Global Biodiversity Framework commitments. Integrative Conservation, 2(3), 140–155. 10.1002/inc3.24

Webber, W., Moffat, A., & Zobel, J. (2010). A similarity measure for indefinite rankings. ACM Transactions on Information Systems, 28(4), 1–38. 10.1145/1852102.1852106

Xu, W., Luo, D., Peterson, K., Zhao, Y., Yu, Y., Ye, Z., Sun, J., Yan, K., & Wang, T. (2025). Advancements in ecological niche models for forest adaptation to climate change: A comprehensive review. Biological Reviews, 100(4), 1754–1781. 10.1111/brv.70023

Zhang, Y., Wang, X., Lian, X., Li, S., Li, Y., Chen, C., & Piao, S. (2024). Asymmetric impacts of forest gain and loss on tropical land surface temperature. Nature Geoscience, 17(5), 426–432. 10.1038/s41561-024-01423-3

Zizka, A., Silvestro, D., Andermann, T., Azevedo, J., Duarte Ritter, C., Edler, D., Farooq, H., Herdean, A., Ariza, M., Scharn, R., Svantesson, S., Wengström, N., Zizka, V., & Antonelli, A. (2019). CoordinateCleaner: Standardized cleaning of occurrence records from biological collection databases. Methods in Ecology and Evolution, 10(5), 744–751. 10.1111/2041-210X.13152

Zou, Y., Crowther, T. W., Smith, G. R., Ma, H., Mo, L., Bialic-Murphy, L., Potapov, P., Gawecka, K. A., Xu, C., Negret, P. J., Lauber, T., Wu, Z., Rebindaine, D., & Zohner, C. M. (2025). Fragmentation increased in over half of global forests from 2000 to 2020. Science, 389(6765), 1151–1156. 10.1126/science.adr6450

Zurell, D., Fritz, S. A., Rönnfeldt, A., & Steinbauer, M. J. (2023). Predicting extinctions with species distribution models. Cambridge Prisms: Extinction, 1, e8. 10.1017/ext.2023.5

